# Two tales of one neural link predict blind individual’s Braille reading proficiency

**DOI:** 10.1101/2021.04.12.439380

**Authors:** Ruxue Wang, Jiangtao Gong, Chenying Zhao, Yingqing Xu, Bo Hong

## Abstract

Natural Braille reading, a demanding cognitive skill, poses a huge challenge for the brain network of the blind. Here, with behavioral measurement and functional MRI imaging data, we pinpointed the neural pathway and investigated the neural mechanisms of individual differences in Braille reading in late blindness. Using resting state fMRI, we identified a distinct neural link between the higher-tier ‘visual’ cortex—the lateral occipital cortex (LOC), and the inferior frontal cortex (IFC) in the late blind brain, which is significantly stronger than sighted controls. Individual Braille reading proficiency positively correlated with the left-lateralized LOC-IFC functional connectivity. In a natural Braille reading task, we found an enhanced bidirectional information flow with a stronger top-down modulation of the IFC-to-LOC effective connectivity. Greater top-down modulation contributed to higher Braille reading proficiency via a broader area of task-engaged LOC. Together, we established a model to predict Braille reading proficiency, considering both functional and effective connectivity of the LOC-IFC pathway. This ‘two-tale’ model suggests that developing the underpinning neural circuit and the top-down cognitive strategy contributes uniquely to superior Braille reading performance.

**SIGNIFICANCE STATEMENT:** For late blind humans, one of the most challenging cognitive skills is natural Braille reading. However, little is known about the neural mechanisms of significant differences in individual Braille reading performance. Using functional imaging data, we identified a distinct neural link between the left lateral occipital cortex (LOC) and the left inferior frontal cortex (IFC) for natural Braille reading in the late blind brain. To better predict individual Braille reading proficiency, we proposed a linear model with two variables of the LOC-IFC link: the resting-state functional connectivity and the task-engaged top-down effective connectivity. These findings suggest that developing the underpinning neural pathway and the top-down cognitive strategy contributes uniquely to superior Braille reading performance.

## Introduction

For blind humans, one of the most demanding cognitive skills is natural Braille reading, which makes a big difference to their social well-being (Ryles, 1996, 2000; Schroeder, 1996; Marshall and Moys, 2020). They decode the tactile dots information into meaningful patterns with semantic properties and ultimately achieve content reading (Sadato, 2005). Learning Braille reading poses a critical challenge to the blind brain, especially to late blind people. Unlike congenital blindness, late blind individuals who lost sight several years after birth may have experienced traditional print early in life and get exposed to Braille only later. Such a huge gap between visual and tactile reading requires more effort for them to learn Braille reading (Trent and Truan, 1997). Moreover, not all blind learners excel at their Braille reading; in fact, they exhibit wide disparities in Braille reading proficiency (Coppins and Barlow-Brown, 2006). However, the neural mechanisms of individual differences in natural Braille reading performance in late blindness remain largely unknown.

Previous studies have reported the recruitment of the somatosensory cortex, the higher-order frontal language system, and the ‘visual’ system during Braille reading in the blind, including late blind humans (Pascual-Leone et al., 2005; Sadato, 2005). This ‘visual’ system stretched from the primary ‘visual’ cortex (V1) to the higher-tier ‘visual’ cortex—the lateral occipital cortex (LOC), which approximately included the consistent region as the typical visual word form area in the sighted (Reich et al., 2011; Kim et al., 2017). Moreover, the most frequently found functional reorganization of the blind brain was the functional connectivity between the ‘visual’ cortex and the inferior frontal cortex (IFC) (Liu et al., 2007; Sabbah et al., 2016; Bedny, 2017), which hinted its role in higher-cognitive functions, including Braille reading. However, previous studies leave open three important questions. First, for people with a late acquisition of blindness and Braille reading, what is the specific ‘visual’ region playing the essential role in bridging the tactile and language-related processing in natural Braille reading? Furthermore, employing either only Braille letter (Cohen et al., 1997) or word recognition (Sadato et al., 1998; Katarzyna Rączy et al., 2019), prior findings have limited functional inference of the dynamic neural interactions under real-life, natural Braille reading scenario. Thus, how does the information flow between the specific ‘visual’ region and the IFC during natural Braille reading? Finally, how can such neural interactions determine the Braille reading proficiency of late blind individuals? In other cognitive behaviors, individual performance has been suggested to correlate with the specific neural connectivity (Jung et al., 2018; Ren et al., 2018). Therefore, we hypothesized that the strength of the distinct resting-state functional connectivity and the task-engaged effective connectivity between the ‘visual’ cortex and the frontal cortex would jointly predict Braille reading proficiency of late blind individuals.

To answer these questions, we tested individual differences in Braille reading proficiency and investigated brain signatures from both resting and task-engaged functional magnetic resonance imaging (fMRI) acquisition in late blindness. We used natural Braille articles in this study to resemble real-life natural Braille reading (Figure 1). We first examined whether the V1 or the LOC in the late blind brain developed resting-state functional connectivity with the IFC compared to sighted controls. We then employed dynamic causal modeling (DCM) to infer their effective connectivity in a natural Braille reading task. To test our hypothesis, we examined how individual Braille reading proficiency was linked with the distinct ‘visual’-frontal neural connectivity. Finally, a neural predictive model was established to further reveal the neural mechanisms of individual differences in Braille reading proficiency.

**Figure 1.**
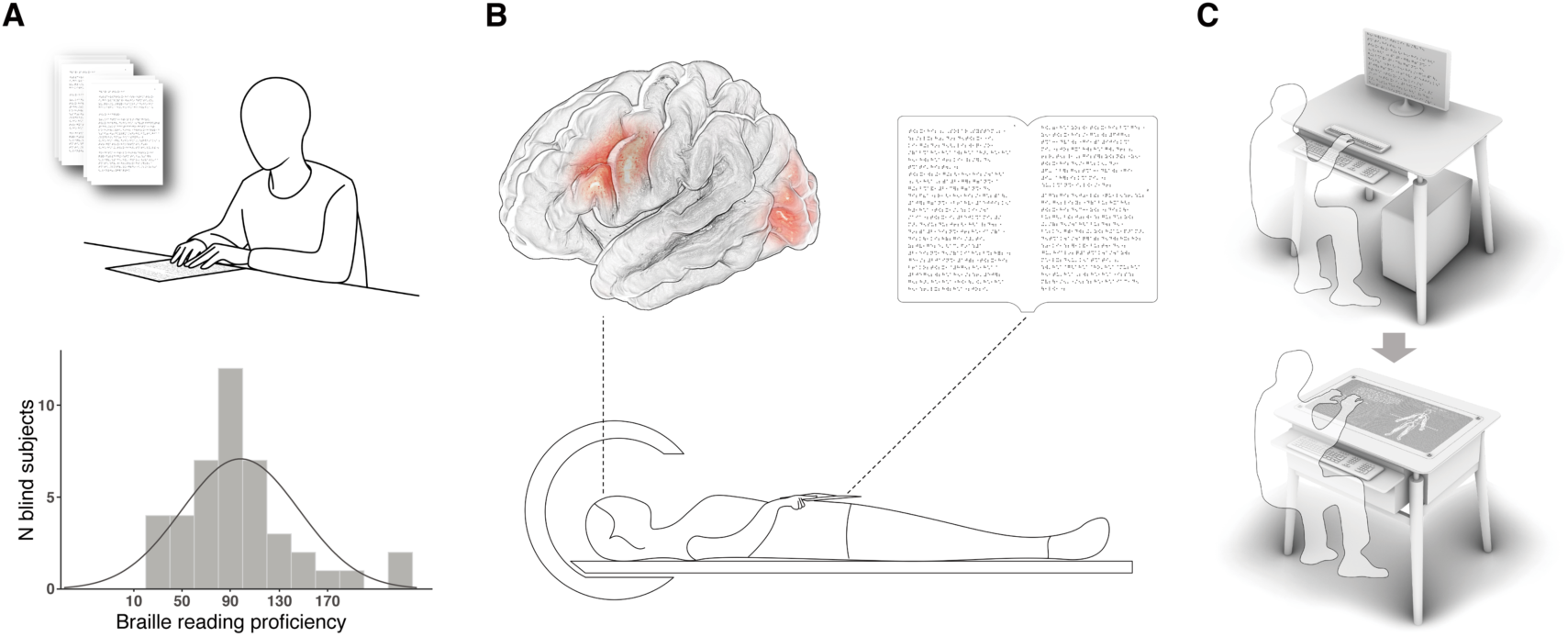
Schematic overview of the study. (A) A natural Braille reading test showed a wide difference in Braille reading proficiency of blind individuals. (B) Both resting-state signals and dynamic task-related fMRI data were assessed. The whole-brain resting-state functional connectivity of two visual brain regions (V1 and LOC), respectively, especially with the frontal cortex, was compared between late blindness and sighted groups. The task fMRI paradigm was designed to simulate real-life Braille reading using natural Braille articles. (C) The current study might inspire the design change from a traditional Braille display (upper panel) with limited single-line Braille words each time to a natural Braille reading device (lower panel) with a continuous Braille article display for blind individuals.

## Materials and Methods

### Experimental subjects

A total of 39 blind subjects (mean age ± s.d. = 21.78 ± 1.96 years; 15 females) participated in the Braille reading performance test. They were all Braille readers. A subgroup of 16 right-handed blind subjects with late-onset acquired blindness (Table 1, mean age ± s.d. = 22.43 ± 1.86 years; mean age at blindness onset ± s.d. = 9.42 ± 4.32 years; 8 females) agreed to participate in the fMRI scan. Fifteen education-matched sighted controls also participated in the fMRI scan. One sighted subject with excessive head motion (mean framewise displacement [FD (Power et al., 2012); > 2 SD from the group mean]) during fMRI acquisition was excluded from further analyses, leaving 14 sighted subjects (mean age ± s.d. = 21.73 ± 1.07 years; 7 females). The two groups were matched in age (two-sample *T*-test, *n* = 16 in the late blind group and *n* = 14 in the sighted group, *T*-value = 1.248, 95% CI = [-0.452 1.862], DF = 28, *p* = 0.222, Cohen’s *d* = 0.457, two-tailed). Sighted subjects had normal or corrected-to-normal vision and had never learned Braille. All subjects had no history of neurological disorders. The study was approved by the Institutional Review Board of Tsinghua University, and informed consent was obtained from each participant.

**Table 1.**
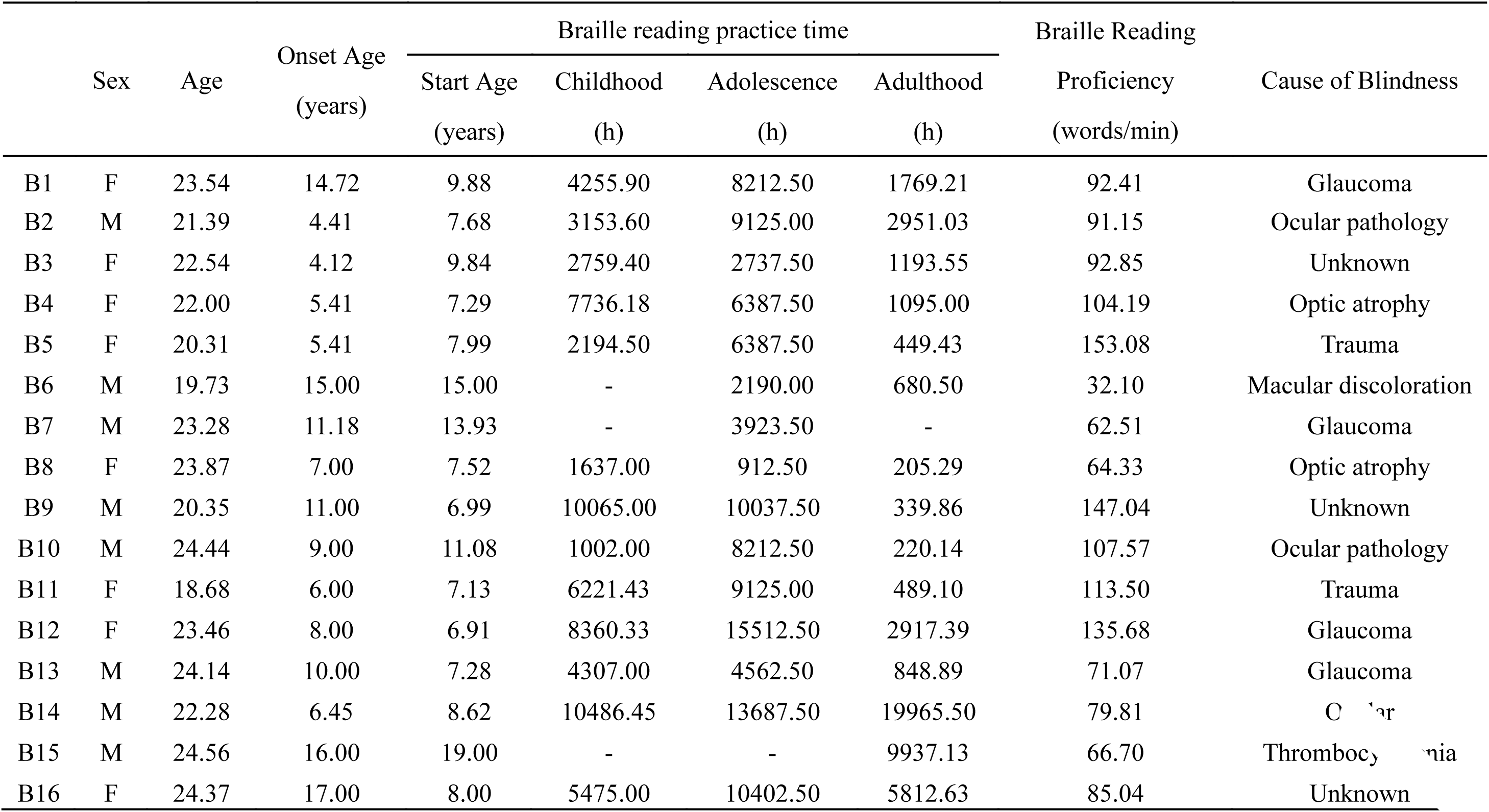
Characteristic of blind participants of the fMRI scan.

|  | Sex | Age | Onset Age<br>(years) | Braille reading practice time |  |  |  | Braille Reading |  |
| --- | --- | --- | --- | --- | --- | --- | --- | --- | --- |
|  |  |  |  | Start Age<br>(years) | Childhood<br>(h) | Adolescence<br>(h) | Adulthood<br>(h) | Proficiency<br>(words/min) | Cause of Blindness |
| B1 | F | 23.54 | 14.72 | 9.88 | 4255.90 | 8212.50 | 1769.21 | 92.41 | Glaucoma |
| B2 | M | 21.39 | 4.41 | 7.68 | 3153.60 | 9125.00 | 2951.03 | 91.15 | Ocular pathology |
| B3 | F | 22.54 | 4.12 | 9.84 | 2759.40 | 2737.50 | 1193.55 | 92.85 | Unknown |
| B4 | F | 22.00 | 5.41 | 7.29 | 7736.18 | 6387.50 | 1095.00 | 104.19 | Optic atrophy |
| B5 | F | 20.31 | 5.41 | 7.99 | 2194.50 | 6387.50 | 449.43 | 153.08 | Trauma |
| B6 | M | 19.73 | 15.00 | 15.00 | - | 2190.00 | 680.50 | 32.10 | Macular discoloration |
| B7 | M | 23.28 | 11.18 | 13.93 | - | 3923.50 | - | 62.51 | Glaucoma |
| B8 | F | 23.87 | 7.00 | 7.52 | 1637.00 | 912.50 | 205.29 | 64.33 | Optic atrophy |
| B9 | M | 20.35 | 11.00 | 6.99 | 10065.00 | 10037.50 | 339.86 | 147.04 | Unknown |
| B10 | M | 24.44 | 9.00 | 11.08 | 1002.00 | 8212.50 | 220.14 | 107.57 | Ocular pathology |
| B11 | F | 18.68 | 6.00 | 7.13 | 6221.43 | 9125.00 | 489.10 | 113.50 | Trauma |
| B12 | F | 23.46 | 8.00 | 6.91 | 8360.33 | 15512.50 | 2917.39 | 135.68 | Glaucoma |
| B13 | M | 24.14 | 10.00 | 7.28 | 4307.00 | 4562.50 | 848.89 | 71.07 | Glaucoma |
| B14 | M | 22.28 | 6.45 | 8.62 | 10486.45 | 13687.50 | 19965.50 | 79.81 | Ocular |
| B15 | M | 24.56 | 16.00 | 19.00 | - | - | 9937.13 | 66.70 | Thrombocytopenia |
| B16 | F | 24.37 | 17.00 | 8.00 | 5475.00 | 10402.50 | 5812.63 | 85.04 | Unknown |

### Acquisition of biographical data on Braille reading practice

According to previous studies (Bengtsson et al., 2005; Liu et al., 2007), we obtained the number of hours of practicing Braille reading for each blind subject during childhood (from the start of practicing until 11 years), adolescence (12–16 years) and adulthood (17 years until the time of the magnetic resonance scan). These values were calculated from biographical data collected from all blind participants on the key events from the beginning of Braille reading training until the scan and on their self-estimated number of hours of practicing within the periods between these key time points (Table 1).

### Braille reading performance test

Previous Braille reading studies have demonstrated that passage length and content validity might influence the reliability and validity of the behavioral assessment (Mason, 2012). The content validity of Braille materials concerns the readability and difficulty for subjects. Given that all of our blind subjects had passed the special college entrance examination for blind individuals, we selected two natural Braille articles at the high school level, slightly below their current levels. Each natural Braille article consisted of approximately 500 Braille words. All reading materials were proofread by an experienced Braille reader and were printed by a Braille printer from VIEWPLUS (Viewplus.com) on Braille paper, measuring 27.94 cm in length and 24.89 cm in width. The task was to read the two natural Braille articles aloud as accurately and quickly as possible. The reading strategy was not specified. We tested individual Braille reading proficiency by measuring the number of Braille words read correctly per minute on average. Individual Braille reading accuracy was also assessed by calculating the proportion of Braille words read correctly to all the words in the two Braille articles. Blind individuals with a reading accuracy reaching 99.5% were recruited for this study to ensure content comprehension.

### Functional MRI parameters

All MR images were acquired from a Philip Achieva 3.0 Tesla TX MR scanner with a 32-channel head coil. The participants laid supine with their heads snugly fixed with straps and foam pads to minimize head movement. Structural images were acquired using a sagittal magnetization-prepared rapid gradient-echo T1-weighted sequence (TR = 7.6 ms, TE = 3.7 ms, flip angle = 8°, voxel size = 1 × 1 × 1 mm, FOV = 256, slices = 180). Before the task session, participants underwent two resting-state fMRI scans (8 min per scan) using an echo-planar imaging sequence (TR = 2000 ms, TE = 30 ms, flip angle = 90°, voxel size = 3.5 × 3.5 × 3.5 mm, FOV = 224, slices = 35). During the resting-state fMRI scan, the participants were instructed to stay awake. Functional images for the natural Braille reading task were also acquired using an echo-planar imaging sequence (TR = 3000 ms, TE = 35 ms, flip angle = 90°, voxel size = 3 × 3 × 3 mm, FOV = 220, slices = 47). All subjects were asked to keep their eyes closed during the scans.

### FMRI task paradigm

A natural Braille reading task with ten blocks was used in this study (Supplementary Figure 4). In each block, blind subjects were instructed to silently read the page of the Braille article and the next blank page (control: blank), mainly using the right index finger at a constant speed, according to the audio cue of “Begin” or “Turn the page” played in an earphone. Blind individuals were asked to thoroughly read and comprehend the Braille article during the fMRI scan. All subjects were instructed to carefully touch and perceive the blank page line by line as if they were reading the Braille page. Each audio cue lasted for 1 s. We used natural Braille articles to simulate real-life Braille reading conditions. Blank pages were printed with the same kind and size of paper as Braille pages.

### Resting-state fMRI data preprocessing and analysis

Structural data were processed and reconstructed to individual cortical surface using FreeSurfer (http://surfer.nmr.mgh.harvard.edu). Individual resting-state data were preprocessed using the following procedures: (i) discarding of the first four volumes; (ii) slice-timing correction; (iii) rigid body correction for head motion with the AFNI package (Cox, 1996); (iv) functional-anatomical image co-registration; (v) bandpass temporal filtering (0.01-0.08 Hz); and (vi) regressing out of nuisance covariates to control for physiological effects and head motion (6 head motion parameters and 2 regressors corresponding to the ventricular signal and white matter signal). One blind subject (B14) with excessive head motion (mean FD (Power et al., 2012) > 2 SD from the group mean) during resting-state fMRI acquisition was excluded, leaving 15 blind subjects for resting-state analyses. The preprocessed fMRI data of each subject were first aligned to the reconstructed individual cortical surface in each hemisphere. Then, a 6-mm full-width half-maximum (FWHM) Gaussian smoothing kernel was applied to the fMRI data in the surface space. The smoothed data were then down-sampled to the FreeSurfer template with 2,562 vertices in each hemisphere using the mri_surf2surf function in FreeSurfer. The fMRI data of two resting-state scans were concatenated for further analyses to improve reliability.

The primary visual cortex (V1) was defined as the combination of the occipital pole and calcarine sulcus in FreeSurfer’s automatic anatomical parcellation (aparc2009) (Destrieux et al., 2010), consistent with the location of V1 in previous studies (Bedny et al., 2011; Watkins et al., 2012; Butt et al., 2013). In accord with the seed region in prior functional connectivity analyses of blind and sighted individuals (Bedny et al., 2011; Watkins et al., 2012; Abboud and Cohen, 2019), we used the lateral occipital cortex as a higher-tier visual seed located in the anatomical occipital region according to the FreeSurfer aparc2009 atlas (Destrieux et al., 2010) (encompassing the middle occipital gyrus and sulcus, anterior occipital sulcus, inferior occipital gyrus and sulcus, and lateral occipital-temporal sulcus). The V1 and the LOC were used as the seeds for resting-state functional connectivity (rsFC) analyses, respectively.

To obtain each seed’s whole-brain functional connectivity patterns, we computed the correlation between the mean blood oxygen level-dependent (BOLD) signal of the seed and the time series of every other vertex in both hemispheres for each subject. These *r*-maps were next converted into *z*-maps using Fisher’s *r*-to-*z* transformation. All statistical group-level FC pattern analyses were carried out using the Surface-Based Data Processing & Analysis for (Resting-State) Brain Imaging software (DPABISurf 1.1) (Yan et al., 2016). A one-sample *T*-test (*n* = 15, DF = 14 in the blind group and *n* = 14, DF = 13 in the sighted group, one-tailed) was performed for individual FC *z*-maps to generate significant and positive FC maps (vertexwise *p* < 10^-6^, uncorrected, cluster size > 100 mm) of the V1 or LOC seed for each group. Thus, a union mask including identified significant vertices in either the blind or sighted group was created for further FC pattern comparison between the two groups (Wang et al., 2015). Recent works have consistently suggested that the influence of head motion on intrinsic functional connectivity in the analysis of between-group functional connectivity comparison needs to be further scrutinized regardless of the correction one employs at the individual level (Power et al., 2012; Yan et al., 2013). We, therefore, calculated the mean FD (Power et al., 2012), describing the mean absolute displacement of each brain volume relative to the previous volume in translation and rotation in the *x*, *y*, and *z* directions during the resting-state scan for each subject.

With mean FD as the covariate controlling for the head motion effect (Wang et al., 2015), we conducted a two-sample *T*-test (*n* = 15 in the blind group and *n* = 14 in the sighted group, DF = 27, two-tailed) based on individual FC *z*-maps within the union mask to examine how the V1 or LOC FC patterns of the blind group changed compared to the sighted group. Significant vertices were those surviving the multiple comparison correction using non-parametric permutation testing (Winkler et al., 2016) (5,000 permutations) and threshold-free cluster enhancement (Chen et al., 2018) (*p* < 0.01, corrected for both hemispheres, cluster size > 30 mm).

### Task fMRI data preprocessing and analysis

The task fMRI data were preprocessed using the FsFast software package (http://surfer.nmr.mgh.harvard.edu/fswiki/FsFast). The preprocessing procedures included motion correction, slice timing correction, and boundary-based functional-anatomical alignment. Like the resting-state fMRI, the task fMRI data of each subject were registered to the reconstructed individual cortical surface. Next, the fMRI data were smoothed with a Gaussian filter (FWHM = 6 mm) in the surface space. The smoothed data were then down-sampled to the FreeSurfer template with 2,562 vertices in each hemisphere. One blind subject (B6) out of 16 with excessive head motion (mean FD (Power et al., 2012); > 2 SD from the group mean) during the task fMRI scan was excluded from further general linear model (GLM) and DCM analyses.

Functional MRI analyses were carried out using Statistical Parametric Mapping toolbox (SPM12, Welcome Department of Imaging Neuroscience, Institute of Neurology, London). For the first-level individual analysis, functional images were modeled with three regressors (one for the Braille condition, one for the blank condition, and another for the turning page condition which was discarded in the analysis) using the GLM method. Head motion regression was also performed in the analysis. We defined a Braille > blank contrast to reveal activated brain regions (significant *p* < 0.025 for both hemispheres, uncorrected). We did not set an extremely strict threshold here because these GLM analyses aimed to yield brain regions showing task-relevant effects for further DCM analyses. However, one blind subject (B16) failing to exhibit any activated vertices in three of our ROIs above a very liberal threshold (*p* < 0.1, uncorrected) was excluded, leaving 14 blind subjects for group-level GLM analysis and DCM analysis. For group-level GLM analysis, data of all the blind subjects were concatenated and then analyzed by a one-sample *T*-test (*n* = 14 in the blind group, DF = 13, one-tailed). Significant brain activations in the blind group were also revealed by the Braille > blank contrast (Supplementary Figure 4, significant *p* < 0.025 for both hemispheres, uncorrected).

### Dynamic causal modeling (DCM)

All DCM analyses were carried out using SPM12. DCM is a computational framework for explaining the effective connectivity among brain regions that show experimental effects (Friston et al., 2003). With the specified model and priors on the coupling and hemodynamic parameters, posterior probability analysis in DCM determines the most likely coupling parameters at the neuronal level. We focused on the dynamic interactions among the IPS, LOC, V1, and IFC in the left hemisphere that were activated when the blind subjects were reading the natural Braille article. The exact left V1, LOC, and IFC regions in the resting-state analysis and the anatomical region of the IPS (intraparietal sulcus, defined in the FreeSurfer aparc2009 atlas (Destrieux et al., 2010)) were used as anatomical masks for defining ROIs later. Individual ROIs consisted of significant vertices revealed by the Braille > blank contrast (*p* < 0.025 uncorrected, cluster size > 5 vertices) within the corresponding anatomical masks. The time series of these ROIs were then extracted for each subject. We slightly dropped the threshold for two blind subjects (B11: *p* < 0.025 uncorrected; B14: *p* < 0.05 uncorrected, cluster size > 5 vertices) who lacked strong responses in one or more of the ROIs to ensure that significant vertices could be identified (Zeidman et al., 2019).

Typically, there are three types of DCM parameters (Friston et al., 2003): 1) endogenous parameters, reflecting the task-independent effective connectivity among brain regions; 2) driving effects, which quantify how neural responses change due to the task stimuli; and 3) modulatory strengths, the parameters of greatest interest, which account for the changes in specific effective connectivity due to the task-relevant conditions. We constructed bidirectional endogenous connections among the four ROIs in DCM models based on anatomical knowledge and our resting-state functional connectivity results. We used the IPS to model the tactile input for the LOC, consistent with the finding that tactile information from the somatosensory cortex was redirected to the ‘visual’ systems through the poly-sensory intraparietal cortex in blind people (Wittenberg et al., 2004; Sadato, 2005; Fujii et al., 2009). We set the modulatory effect of Braille reading on the connectivity between the left IPS and LOC in accord with a prior study (Fujii et al., 2009). We then investigated the task modulatory effect of Braille reading on the connection between the left LOC and the left IFC. Therefore, we considered that the task-relevant information flow on any direction (bottom-up or top-down) between these two brain regions might be modulated, resulting in four competing models (Figure 4A).

We performed BMS across all blind individuals with random-effects analysis to select the winning model after balancing the explanation of data and the model complexity. The exceedance probability intuitively presented the relative preference across all models (Figure 4B). BMA implements a weighted average of parameters in all competing models where the weights were given by the evidence of each model (Penny et al., 2010). We then employed BMA to estimate the full posterior density on model parameters (Penny et al., 2010). Therefore, the significance of each connectivity parameter could be assessed by the fraction of samples in the posterior density (out of 10,000 sampled data points) that were different from zero (Seghier et al., 2011). Likewise, the significance of the difference in modulatory strengths on one connection versus another connection was assessed by the fraction of samples that were higher for one connection than another (Seghier et al., 2011).

### Statistical analysis on the relationship between resting-state functional connectivity and Braille reading behavior data

To investigate the statistical relationship between the left LOC-IFC neural link and Braille reading behavior data of blind individuals, we calculated the Pearson correlation and Spearman correlation coefficients between their functional connectivity (normalized) and Braille reading proficiency, as well as Braille reading practice time within each age periods respectively. In addition, we also performed partial correlations between the left LOC-IFC functional connectivity and Braille reading behavior data with covariates of the onset age of blindness, the current age, and motion in the resting state scan indexed by mean FD (Jung et al., 2018).

### Statistical analysis on the relationship between top-down effective connectivity and Braille reading proficiency

The Pearson correlation between top-down modulation of the left IFC-to-LOC effective connectivity in the natural Braille reading task and individual Braille reading proficiency tended to approach significance (*n* = 13, *p* = 0.141, *r* = 0.432, Supplementary Figure 5A). The top-down modulation parameter of DCM analysis provided a precise estimation at the neuronal level of how the rate of change of activity in the IFC influences the rate of change in the LOC activity (Friston et al., 2003) in the natural Braille reading task. Hence, to fully investigate the potential relationship between the top-down modulation and individual Braille reading performance, we investigated whether the area of functional LOC could act as a neural mediator of the top-down modulation in support of the natural Braille reading task. In accord with our DCM analysis, the neural activity of the functional LOC in the task was predominantly influenced by the changes of activity in the IFC through the top-down effective connectivity. Accordingly, we first identified individual peak vertex revealed by the Braille > blank contrast as the center of individual functional LOC. Within the anatomical LOC cortex, individual functional LOC also consisted of vertices whose functional connectivity (normalized) with the peak vertex reached the threshold. The area of individual functional LOC was further normalized by calculating the proportion of these vertices in the anatomical LOC. We initially set a threshold value throughout the functional connectivity with the peak vertex in 0.2 intervals and eventually chose the threshold value of 0.8 since only when using this threshold did the area of functional LOC manifest significant correlation with both top-down modulation (Figure 5A) and Braille reading proficiency of blind individuals (Figure 5C). However, the bottom-up modulation of the LOC-to-IFC connectivity was correlated with the area of functional LOC using none of these threshold values.

Mediation analysis was performed using the “mediation” package (Tingley et al., 2014) in R (https://www.r-project.org). To test the significance of the mediation effect, we used the bootstrapping method (simulations: 5000, the bias-corrected and accelerated confidence intervals were estimated then), which was generally considered more advantageous.

### Analysis of linear neural predictive models for individual Braille reading proficiency

Based on our hypothesis that the resting-state functional connectivity (rsFC) and task-specific top-down effective connectivity (tdEC) of the left LOC-IFC neural connection each explained unique individual variance, we first established a ‘two-tale’ model linearly combining these two variables to predict Braille reading proficiency of late blind individuals. Note that in the model, the area of functional LOC was used as the neural surrogate of the tdEC (Figure 5D). In this case, this model is given by:

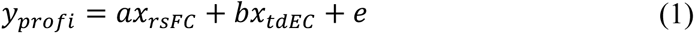

Then, derived from Eq (1), linear models with each of the two brain variables alone, either *x_rsFC_* or *x_tdEC_*, were fitted separately.

We assessed the linear models’ performance in predicting the Braille reading proficiency of a held-out individual using a leave-one-out cross-validation procedure. On each held-out data of this cross-validation procedure, we fitted the model to the training set by minimizing the following cost function:

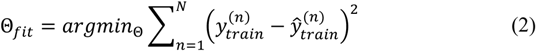

Here, the 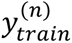 term refers to the measured Braille reading proficiency of the *n-*th blind subject in the training set (*N* = 12), while the 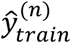 term refers to the *n-*th individual behavior data estimated by each model during training. This minimization was performed using *fmincon* in MATLAB R2018a. We then applied these model parameters to predict the Braille reading proficiency of the held-out test individual. Finally, the MSE and the Pearson correlation between predicted and corresponding actual proficiency for blind individuals from the cross-validation procedure were calculated for each model.

## Results

### Individual differences in Braille reading proficiency

We assessed the Braille reading proficiency of blind individuals using a natural Braille reading behavioral test. Thirty-nine blind participants (mean age ± s.d. = 21.78 ± 1.96 years; 15 females) read two long natural Braille articles (500 Braille words on average) aloud as accurately and quickly as possible. Results showed that the Braille reading proficiency of blind individuals varied considerably (Figure 1A, Braille words correctly read per minute, mean ± s.d. = 97.80 ± 43.31). Sixteen of these participants with late-onset acquired blindness participated in the subsequent functional imaging experiments (Table 1, mean age ± s.d. = 22.43 ± 1.86 years; Braille reading proficiency, mean ± s.d. = 93.69 ± 32.63; 8 females).

### Distinctive functional connectivity between the higher-tier ‘visual’ cortex and the frontal cortex in the late blind brain

Taking the sighted group as the reference, we performed a whole-brain seed-to-vertex-based functional connectivity analysis to test connectivity changes in the late blind group of two visual brain regions, V1 and LOC, for the two hemispheres separately. The whole-brain functional connectivity for V1 was not different between the blind and sighted groups. In both groups, we found significant connections of V1 with other visual areas but not with frontal language areas—the IFC (Figure 2, Supplementary Figure 1, right panel). By contrast, we observed significantly enhanced functional connectivity of the left LOC with the left IFC in the blind group relative to the sighted group (significant difference at *p* < 0.01, corrected for both hemispheres; Figure 2, left panel). Strong connections of the LOC with other visual regions and extensively with dorsal parietal regions were similarly found in both groups (Figure 2, Supplementary Figure 1, left panel). Notably, despite a significant connection between the left LOC seed and the right IFC in the blind group (*p* < 10^-6^, uncorrected), the difference in such connectivity with the sighted group failed to reach a significant level (thresholded at *p* < 0.01, corrected for both hemispheres; Figure 2, left panel). In addition, we also found significantly stronger functional connectivity between the left LOC and the left intraparietal sulcus (IPS) in the blind group than in the sighted group (Supplementary Figure 2).

**Figure 2.**
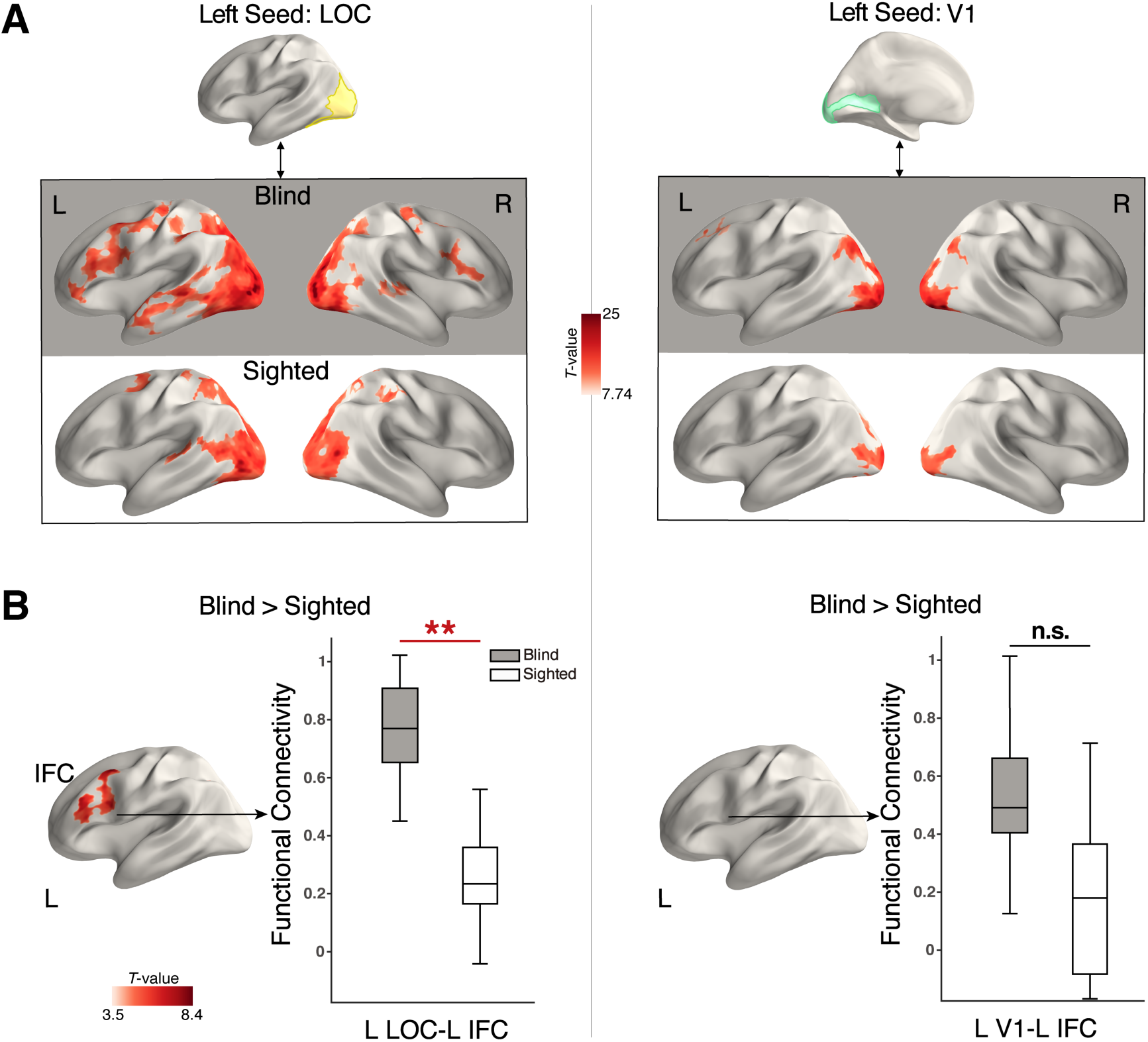
Functional connectivity patterns for the primary ‘visual’ cortex (V1) and the lateral occipital cortex (LOC). (A) Maps show vertices with significantly positive connectivity with the left LOC seed (vertexwise *p* < 10^-6^, uncorrected, cluster size > 100 mm) in the late blind group and sighted group (left panel). Maps show vertices with significant positive connectivity with the left V1 seed (vertexwise *p* < 10^-6^, uncorrected, cluster size > 100 mm) in the late blind group and sighted group (right panel). (B) Differences in functional connectivity patterns for left LOC (left panel) and V1 (right panel) in the late blind group versus the sighted group, respectively. Significant vertices were corrected for multiple comparisons using permutation testing with threshold-free cluster enhancement (*p* < 0.01, corrected for both hemispheres, cluster size > 30 mm). Box plots of the two groups’ connectivity strengths (Fisher *r*-to-*z*-transformed) between the seed and the left IFC cluster (located from the left panel). The horizontal line denotes the median, and the asterisk indicates significance. V1 primary visual cortex (light green), LOC lateral occipital cortex (light yellow), IFC inferior frontal cortex. L left hemisphere, R right hemisphere. Blind, dark color; Sighted, light color. \*\**p* < 0.01, corrected for both hemispheres. n.s. non-significant.

### Left LOC-IFC functional connectivity correlated with Braille reading proficiency of late blind individuals

To test our hypothesis, we examined whether there was an association between the left LOC-IFC functional connectivity and individual levels of Braille reading proficiency. We found that the left LOC-IFC functional connectivity was significantly and positively correlated with individual Braille reading proficiency (Figure 3A left panel, Pearson correlation, *n* = 15, *p* = 0.023, *r* = 0.580; Spearman correlation, *n* = 15, *p* = 0.005, *r* = 0.686). This effect remained significant after controlling for the age of blindness onset, current age, and mean framewise displacement (Power et al., 2012) (motion) in the resting-state scan (partial Pearman correlation, *p* = 0.101, *r* = 0.496; partial Spearman correlation, *p* = 0.011, *r* = 0.704). By contrast, the left V1-IFC functional connectivity was not correlated with the blind individual’s Braille reading proficiency (Figure 3B left panel, Pearson correlation, *n* = 15, *p* = 0.998, *r* = -0.001; Spearman correlation, *n* = 15, *p* = 0.870, *r* = 0.046). Thus, better Braille reading proficiency was associated with greater left LOC-IFC functional connectivity.

**Figure 3.**
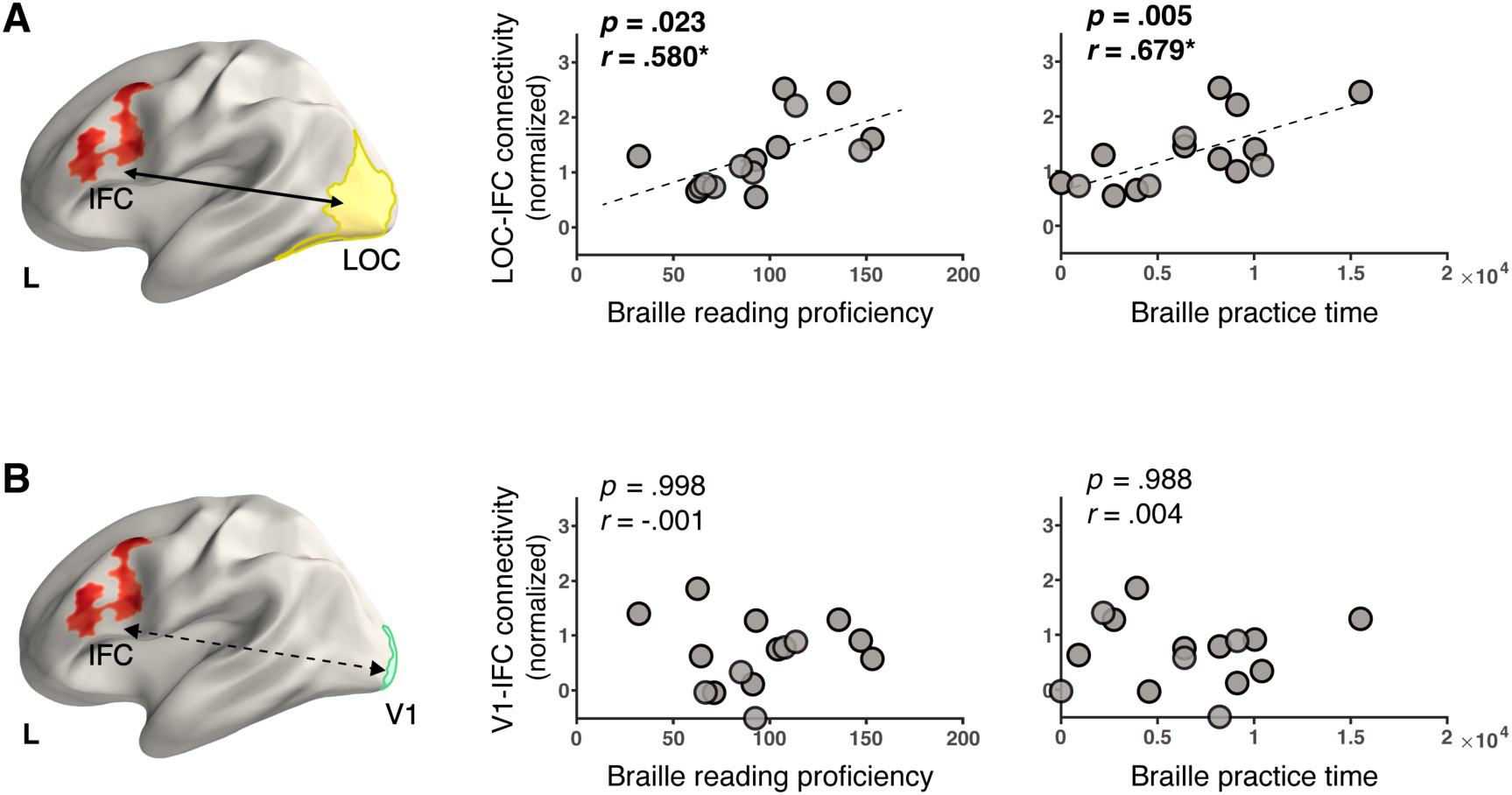
Left LOC-IFC functional connectivity showed significant correlations with individual Braille reading proficiency and practice time. (A) Plots of the left LOC-IFC functional connectivity. Significant and positive correlations were found between individual strengths of the left LOC-IFC functional connectivity and Braille reading proficiency (left panel) and Braille reading practice time in adolescence (in hours, right panel). (B) The connectivity between the left V1 and the left IFC was not correlated with either Braille reading proficiency (left panel) or Braille reading practice time (right panel). L left hemisphere. All *r* and *p*-value pairs reflect Pearson correlation, consistent with the Spearman correlation results.

Next, following previous studies (Liu et al., 2007), we calculated the estimated number of hours of practicing Braille reading for late blind participants during childhood, adolescence, and adulthood (Table 1). We further examined whether the left LOC-IFC functional connectivity was also associated with the individual number of hours of practicing Braille reading. We found that the Braille reading practice time during adolescence was significantly correlated with the left LOC-IFC functional connectivity in blind individuals (Figure 3A right panel, Pearson correlation, *n* = 15, *p* = 0.005, *r* = 0.679; Spearman correlation, *n* = 15, *p* = 0.022, *r* = 0.586; in other age periods, Supplementary Figure 3A). This positive relationship remained significant after controlling for the age of blindness onset, current age, and head motion (Power et al., 2012) (partial Pearson correlation, *p* = 0.027, *r* = 0.634; partial Spearman correlation, *p* = 0.076, *r* = 0.531). By contrast, the left V1-IFC connectivity was not associated with blind individual’s Braille reading practice time (Figure 3B right panel, in adolescence, Pearson correlation, *p* = 0.988, *r* = 0.004; Spearman correlation, *p* = 0.970, *r* = 0.011; in other age periods, Supplementary Figure 3B). Thus, stronger left LOC-IFC functional connectivity was associated with more Braille reading practice.

### Bidirectional LOC-IFC information flow for Braille reading with a preponderance in the top-down modulation

Next, we investigated the direction of the left LOC-IFC information flow and how the information flow was modulated by the Braille reading task, employing dynamic causal modeling (DCM). The blind subjects were asked to silently read pages containing natural Braille articles while undergoing an fMRI scan, with no-Braille blank pages as the control. The Braille > blank contrast elicited bilateral V1, LOC, IPS, and left IFC activations in blind subjects (Supplementary Figure 4). In addition, we constrained our DCM models to the left hemisphere due to the left-lateralized activation of the IFC and the left-lateralized increased LOC-IFC and LOC-IPS functional connections in blind individuals. Therefore, four brain regions, IPS, V1, LOC, and IFC—all frequently implicated in Braille reading (Sadato, 2005; Katarzyna Rączy et al., 2019)—were selected for further DCM analysis.

We specified the model space as shown in Figure 4A. The null model represents the basic structure, including endogenous effective connectivity among regions of interest (ROIs) and the modulation of the IPS-LOC connection confirmed in the previous study (Fujii et al., 2009). We used the IPS to model the tactile input for the LOC because previous studies (Wittenberg et al., 2004; Sadato, 2005; Fujii et al., 2009) agreed that the tactile information from the somatosensory cortex was redirected to the occipital cortex through the poly-sensory intraparietal cortex. To investigate the dynamic interactions between the left LOC and the left IFC in late blind humans, each direction of this effective connectivity can be either modulated or not, hence obtaining four possible models. Random-effects Bayesian model selection (BMS) across blind individuals showed that the bidirectional model largely outperformed other models (Figure 4B, exceedance probability [the probability that a particular model is more likely than any other model] of the bidirectional model = 94.56%).

**Figure 4.**
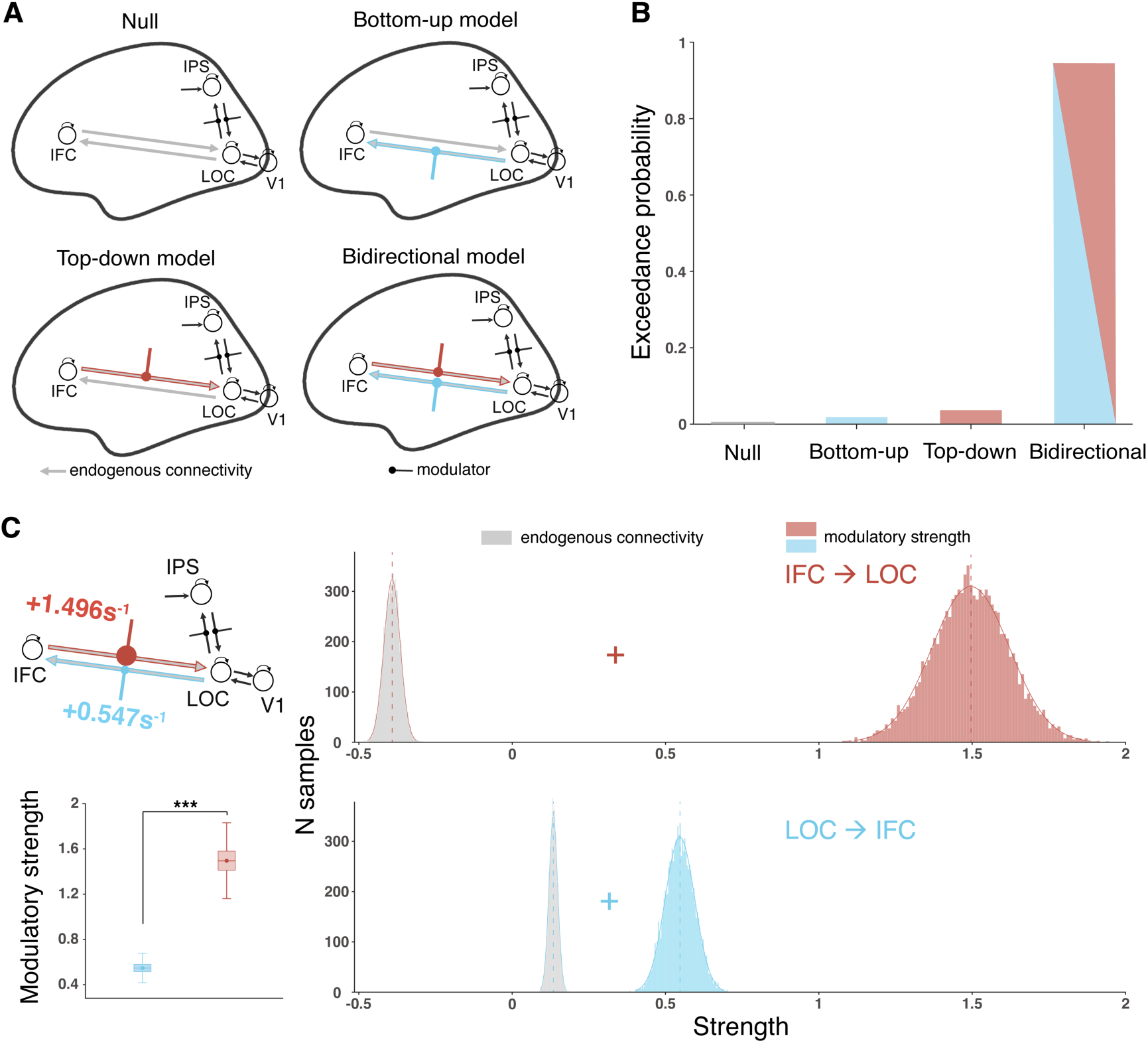
DCM analyses of modulations of the left LOC-IFC effective connectivity for natural Braille reading. (A) Model space specified by adding task modulation to different directions of the left LOC-IFC connection, resulting in four competing models. (B) Exceedance probability for the DCM models. (C) Illustration of the Bayesian model averaging (BMA) result for the late blind group (all significant at a posterior probability threshold of 0.99). Modulators added to the LOC-IFC connection are coded in the size of the filled circles, representing their effect size (left panel). The modulatory strength (in Hz) of Braille reading on the IFC-to-LOC connection was significantly higher than that on the LOC-to-IFC connection (*p* < 0.001). For each parameter (endogenous or modulatory, in Hz) of the top-down IFC-to-LOC connection and the bottom-up LOC-to-IFC connection, the distribution of the 10,000 samples of the posterior densities is provided, respectively. Endogenous connectivity (gray); modulatory strength, red for IFC-to-LOC connection, and cyan for LOC-to-IFC connection. \*\*\**p* < 0.001.

Bayesian model averaging (BMA) over all the competing models showed that the endogenous connectivity and modulatory effect parameters between the LOC and IFC were all significant (Figure 4C, Supplementary Tables 1, 2, posterior probabilities > 0.99). As illustrated in Figure 4C, the LOC-to-IFC endogenous connectivity was positive (0.134 s^-1^), while the IFC-to-LOC connectivity was negative (−0.392 s^-1^). These results indicate that irrespective of the task-relevant modulation, the activity changes of the LOC increased the activity of the IFC but not vice versa. The modulation effects of Braille reading on the left LOC-IFC bidirectional connections were both positive in the late blind group, with a modulation strength of + 0.547 s^-1^ for the bottom-up connection and + 1.496 s^-1^ for the top-down connection (Figure 4C left panel). Notably, the Braille reading task-relevant modulation of the top-down communication from the left IFC to the LOC was significantly stronger than the bottom-up communication from the LOC to the IFC (Figure 4C, *p* < 0.001). Such asymmetrical modulation highlighted the remarkable functional relevance to natural Braille reading of the top-down communication from the left IFC to the left LOC in late blind humans.

### Greater top-down modulation of the left IFC-to-LOC connectivity contributed to higher Braille reading proficiency via a broader area of task-engaged LOC

Significant and positive top-down modulation of the IFC-to-LOC effective connectivity during the natural Braille reading task suggested that at the neuronal level, the information flow from the left IFC increasingly induced the task-specific activity in the left LOC. Accordingly, to measure such neural activity, we identified the individual peak of activation (Supplementary Figure 5C) of the left LOC during natural Braille reading and surrounding vertices with the strongest functional connection to this LOC peak. We referred to such vertices within the anatomical LOC region as the functional LOC of blind individuals. Significant and positive correlations were found between individual top-down modulatory strength and the area of functional LOC (Figure 5A, Pearson correlation, *n* = 13, *p* = 0.005, *r* = 0.730), as well as between the area of individual functional LOC and Braille reading proficiency (Figure 5C, Pearson correlation, *n* = 13, *p* = 0.011, *r* = 0.675). Thus, we ran a mediation analysis to investigate whether the top-down modulation of the left IFC-to-LOC connectivity was associated with individual Braille reading proficiency through the area of functional LOC. The top-down modulation during the task and individual Braille reading proficiency acted as the independent and dependent variables respectively, whereas the area of functional LOC acted as the neural mediator. Results showed that the area of functional LOC significantly and completely mediated the relationship between top-down modulation and Braille reading proficiency (Figure 5D, mediation effect = 9.282, 95% CI = [1.765, 23.870], *p* = 0.043), providing a neural surrogate of the top-down modulation of the left IFC-to-LOC connection in support of late blind individual’s Braille reading proficiency. In contrast, the bottom-up modulation of the left LOC-to-IFC connection was not significantly associated with either the area of functional LOC (Figure 5B, *n* = 13, *p* = 0.644, *r* = 0.142) or individual Braille reading proficiency (Supplementary Figure 5B).

**Figure 5.**
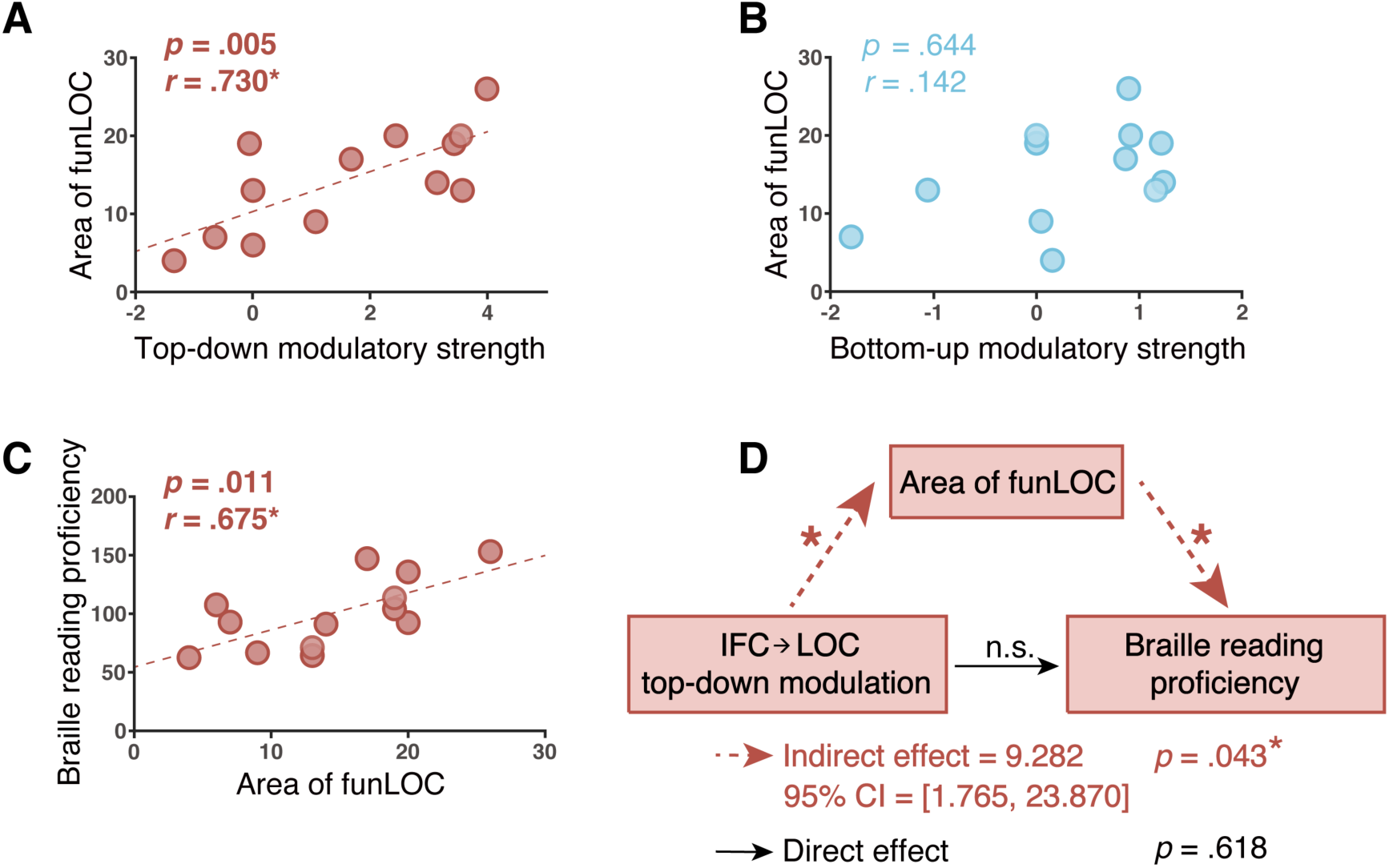
Top-down modulation of the IFC-to-LOC connection facilitated individual Braille reading proficiency via a broader area of functional LOC in the task. (A) Task modulation of the top-down communication from the left IFC-to-LOC (in Hz) was positively correlated with the area of functional LOC (in %) engaging in natural Braille reading. (B) There was no significant relationship between task modulation of the bottom-up communication from the left LOC to IFC (in Hz) and the area of functional LOC (in %). (C) The area of functional LOC engaging in the natural Braille reading task was positively correlated with individual Braille reading proficiency. All *r* and *p*-value pairs reflect Pearson correlation. (D) Mediation analysis showed a significant and complete mediation effect, suggesting that the top-down modulation was significantly associated with enhanced Braille reading proficiency through the increase of the area of functional LOC. FunLOC - functional LOC.

### A ‘two-tale’ model with the LOC-IFC neural link better predicts individual Braille reading proficiency

Finally, we established neural predictive models to further understand the neural mechanisms of individual variance in Braille reading proficiency. We imagined a linear ‘two-tale’ model with two neurofunctional predictors that the resting-state functional connectivity (rsFC) and task-specific top-down effective connectivity (tdEC) of the left LOC-IFC neural link each explained independent variance in individual Braille reading proficiency in late blindness. Here we used the area of functional LOC (Figure 5D) as the neural surrogate of the top-down modulation of the left IFC-to-LOC effective connectivity during the natural Braille reading task. We also ran linear models with each of these two measures of the left LOC-IFC neural connection separately to predict Braille reading proficiency. We assessed the models’ performance for held-out data using a leave-one-out cross-validation procedure. Cross-validation results showed that the ‘two-tale’ model outperformed the other two models in terms of both MSE (Figure 6B, Table 2, mean ± s.e.: ‘two-tale’ model = 457.619 ± 158.048; Model with only rsFC = 661.208 ± 245.886; Model with only tdEC = 651.066 ± 174.217) and *R*^2^ (Table 2, Model 1 = 0.490; Model 2 = 0.285; Model 3 = 0.279) between the predicted proficiency and the held-out actual proficiency. Notably, the coefficients on these two neural measures did not change dramatically, and the total variance explained was near additive when considering these two variables separately and together in different models (Table 2). These results reinforced that for the left LOC-IFC neural link of late blind humans, the rsFC and the tdEC strength each significantly contributed to unique individual variance in Braille reading proficiency in distinct ways.

**Figure 6.**
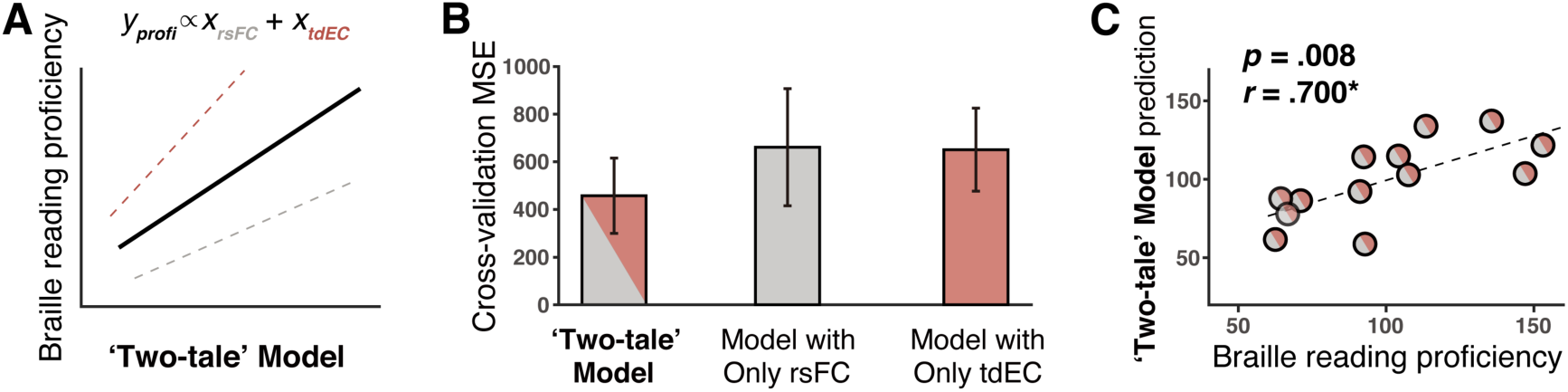
Cross-validation performance of models in predicting Braille reading proficiency of late blind individuals. (A) Schematic representation of the ‘two-tale’ model in predicting Braille reading proficiency of late blind individuals. The resting-state functional connectivity and the task-specific top-down effective connectivity of the left LOC-IFC neural link contributed to unique variance in individual Braille reading proficiency. According to the mediation effect (Figure 5D), the area of functional LOC engaging in the natural Braille reading task was used as the neural surrogate of the top-down effective connectivity from the left IFC-to-LOC here in the model. (B) Leave-one-out cross-validation analysis showed that the ‘two-tale’ model outperformed two other models with a single neural predictor, either the functional or top-down effective connectivity of the left LOC-IFC neural link, in predicting a blind individual’s Braille reading proficiency. (C) A significant and positive correlation was observed between the predicted Braille reading proficiency of the ‘two-tale’ model and the held-out test proficiency. The *r* and *p*-value pair reflect the Pearson correlation. rsFC - resting-state functional connectivity between the left LOC and the left IFC, tdEC - top-down modulation of the left IFC-to-LOC effective connectivity in the natural Braille reading task, profi - individual Braille reading proficiency, MSE - mean squared

**Table 2.** Summary of cross-validation results of linear neural predictive models for Braille reading proficiency of late blind individuals.

|  | ‘Two-tale’<br>Model | Model with<br>Only rsFC | Model with<br>Only tdEC |
| --- | --- | --- | --- |
| rsFC | 21.265 | 29.946 | - |
| tdEC | 229.399 | - | 317.238 |
| constant | 38.775 | 60.348 | 54.455 |
| MSE | 457.619 | 661.208 | 651.066 |
| $R^2$ | 0.490 | 0.285 | 0.279 |

Data are given as unstandardized mean coefficients of the fitted models of the leave-one-out cross-validation procedure. rsFC resting-state functional connectivity between the left LOC and the left IFC, tdEC top-down modulation of the left IFC-to-LOC effective connectivity, which contributed to individual Braille reading proficiency in the models via a neural mediator—the area of functional LOC engaging in the natural Braille reading task, MSE mean squared error between the predicted proficiency and the held-out actual proficiency, $R^2$ $R$ -square between the predicted proficiency and the held-out actual proficiency.

## Discussion

This study linked resting-state functional connectivity and task-engaged effective connectivity from functional imaging data together to natural Braille reading behavior in late blindness. The remarkable resting-state functional connectivity between the higher-tier ‘visual’ cortex—the LOC, but not the V1, and the IFC in the late blind brain, as well as its significant relationship with individual Braille reading proficiency, concurred in suggesting the indispensable role of such functional connectivity in bridging Braille orthographic and language processing of natural Braille reading. In a natural Braille reading task, DCM analyses revealed increased bidirectional information flow with a preponderance in the top-down modulation of the left IFC-to-LOC effective connectivity. Further analyses suggested greater top-down modulation of the left IFC-to-LOC effective connectivity contributed to higher Braille reading proficiency through a mediation effect. Crucially, to fully reveal the neural mechanisms of individual variance in Braille reading performance, we established a neural predictive model with two variables of the LOC-IFC link: the functional connectivity and the top-down effective connectivity. This ‘two-tale’ model suggests that developing the underpinning neural circuit and the top-down cognitive strategy contributes uniquely to superior Braille reading performance. This work potentially provides a general framework for understanding crossmodal neuroplasticity during learning.

The crossmodal recruitment of the typical visual cortex for nonvisual higher cognitions in blind individuals, including Braille reading, has been frequently observed (Pavani and Röder, 2012; Renier et al., 2014; Bedny, 2017; Heimler and Amedi, 2020). A particular concern regarding this plasticity is whether there is a reversed hierarchy of functional organization across ‘visual’ regions or not (Amedi et al., 2003; Büchel, 2003; Burton, 2003; Watkins et al., 2012). Studies by Amedi et al. (2003) first proposed a reversal of the normal visual hierarchy in congenitally blind individuals by showing that compared to higher-tier ‘visual’ regions, early V1 preferentially responded to more ‘abstract’ cognitive tasks (Amedi et al., 2003; Büchel, 2003). These findings are in accord with studies showing functional connectivity of blind individuals between the V1 and frontal language areas at rest (Butt et al., 2013; Striem-Amit et al., 2015). Some studies, on the other hand, did not support such reversed hierarchy by providing evidence of task activations (Röder et al., 2002; Burton, 2003; Noppeney, 2003) and resting-state functional connectivity (Bedny et al., 2011; Watkins et al., 2012) of the primary and higher-tier ‘visual’ regions. For the late blind who lost their sight several years after birth, a normal hierarchy is more possible since late blind humans might have initially developed their neural circuits as normally sighted people (Röder et al., 2021). Here in our study, the functional connectivity between higher-tier ‘visual’ region LOC and the frontal cortex (Figure 2) and its behavioral significance for natural Braille reading (Figure 3) indicates a normal hierarchy. Notably, the anatomical LOC area in this study, approximately encompassed the anatomical location of the visual word form area (VWFA) as in sighted people. Studies showed the ‘VWFA’ activation of blind humans not only in tactile Braille reading but also in functions such as language processing (Reich et al., 2011; Kim et al., 2017; Katarzyna Rączy et al., 2019), which is similar to the functional profile of the broader LOC cortex (Noppeney, 2003; Bedny et al., 2011; Watkins et al., 2012; Bedny, 2017). In line with these findings, our data suggest that the left LOC in late blind humans might incorporate the VWFA-like function and extensively develop as a versatile crossmodal region, bridging Braille orthographic and language processing in natural Braille reading via the distinctive neural nexus of the left LOC-IFC. In line with the connectivity-based plasticity theory (Schepers et al., 2012; Bedny, 2017), the development of such neural connectivity in late blind humans likely depends on the prior connectivity between the VWFA and frontal language areas as normally sighted people.

Our DCM results provide direct evidence for the direction of information flow on the left LOC-IFC neural interaction of late blind humans during a natural Braille reading task. In line with the Hebbian theory (Hebb, 1950), the winning model suggests that natural Braille reading strongly modulates and strengthens the bidirectional LOC-IFC connection (Figures 4A, 4B). This result mirrors the robust resting-state functional connectivity in late blind humans (Figure 2A), as well as its positive relationship with Braille reading practice time (Figure 3A), although the preferential practice time might depend on the age of blindness onset (Liu et al., 2007). However, a closer look at the winning model suggests that such neural connection does not simply convey balanced reciprocal interactions but predominantly delivers the task-specific top-down information from the higher cognitive cortex—the left IFC (Figure 4C). This top-down influence guarantees the active prediction and perception of the tones and semantic meanings through the use of contextual cues (Trent and Truan, 1997; Friston, 2010). Importantly, we found that the task modulation of top-down connectivity from the left IFC to the left LOC was strongly and positively associated with individual Braille reading proficiency through a mediator, the area of task-engaged functional LOC (Figure 5D). Our findings reinforced the significant contribution of the top-down influence to natural Braille reading.

Furthermore, why the ‘visual’ cortex engages more actively in various crossmodal tasks in blind humans, including late blind humans, than in sighted controls (Matuszewski et al., 2021) is still debated (Röder et al., 2021). Using natural Braille reading as a model, the current study provides empirical evidence for the hypothesis that ‘visual’ cortex activity in late blindness predominantly arises from the top-down connectivity from the higher-order multisensory or supramodal cortex (Collignon et al., 2013; Bedny, 2017; Röder et al., 2021). This neural mechanism is essential for the rapid acquisition of new skills because it might be relatively difficult for late blind humans to extract crossmodal characteristics exclusively via bottom-up stimulus input, with their visual deprivation occurring after the pruning of exuberant crossmodal connectivity during the typical early development (Batardière et al., 2002; Röder et al., 2021). Such a top-down controlled learning mechanism in late blind humans is also compatible with the typical adult learning characteristics demonstrated by human and non-human animal studies (Keuroghlian and Knudsen, 2007; Rohlf et al., 2017).

To our knowledge, this is the first study to date to establish the neural predictive model for the natural Braille reading performance of late blind individuals. Our model’s two neurofunctional predictors regarding the left LOC-IFC neural interactions arguably happen by distinct experience-dependent brain plasticity in late blindness. The functional connectivity predictor at the resting state might stem from prolonged Braille reading practicing of late blind humans on a much slower timescale (likely at months to years), which is fundamental to high-level Braille reading performance. In line with previous studies (Bengtsson et al., 2005; Scholz et al., 2009), we postulate that increased myelination resulted from constant neural activity along the fiber tract of the inferior fronto-occipito fasciculus (Forkel et al., 2014) during training, is a potential mechanism underlying the facilitated communications between neural ensembles of the LOC and the IFC. Previous work in animals (Pan et al., 2020) also showed that lifelong learning-induced myelin plasticity affected neuronal responses and subsequent behavior. On the other hand, the effective connectivity predictor in our model represents the effect of selectively implementing a top-down cognitive strategy such as active prediction in every natural Braille reading task on a faster timescale (likely at milliseconds to seconds). Consistent with previous studies (Citri and Malenka, 2008), we speculate that the burst of activity caused by the top-down influence of the IFC largely increases the probability of neurotransmitter release to neurons of the LOC. Such metabolically and behaviorally efficient synaptic plasticity (Mongillo et al., 2008) boosts the information transmission of synapses (Citri and Malenka, 2008) and the LOC activity and, therefore, the behavior performance. If further refined and validated in a larger cohort, this brain-behavior model might be not only beneficial for more efficient acquisition of Braille reading skills in blind individuals, but also instructive for more general learning practices. Future studies are needed to provide detailed neurobiological evidence to explicitly elucidate the mechanism of each neurofunctional predictor.

Unlike other studies using Braille letters or words, the large-scale Braille articles used in our study made it possible to observe task-specific top-down communication in blind humans during natural Braille reading. The ‘two-tale’ model proposed in the current study (Figure 6) suggests that for blind Braille readers to make better use of this top-down effective connectivity during natural Braille reading, a promising strategy might be to make the reading context more available. Compared with traditional refreshable Braille displays offering only a single line of Braille words each time (Ren et al., 2008), emerging technology is the large-print Braille computer (Völkel et al., 2008; Jiao et al., 2018), which enables continuous and large-scale natural Braille context display (Figure 1C). This new type of Braille reading device arguably allows for active understanding and prediction based on contextual information during natural Braille reading, boosting the top-down information flow and thus promoting Braille reading proficiency of blind individuals.

## Acknowledgements

This work is supported by the National Science Foundation of China [grant number NSFC 61621136008 to B.H.] and the National Key R&D Program of China [grant number 2017YFA0205904 B.H.] We thank Brigitte Röder at the University of Hamburg, Ning Guo, and Fang Cai at Tsinghua University for comments on the manuscript.

## Data and code availability

All data reported in this paper will be available by the corresponding author upon request.

All original code has been deposited at Github and is publicly available as of the date of publication.

## References

Abboud S, Cohen L (2019) Distinctive Interaction Between Cognitive Networks and the Visual Cortex in Early Blind Individuals. Cerebral cortex (New York, NY : 1991) 29:4725–4742.

Amedi A, Raz N, Pianka P, Malach R, Zohary E (2003) Early “visual” cortex activation correlates with superior verbal memory performance in the blind. Nature Neuroscience 6:758–766.

Batardière A, Barone P, Knoblauch K, Giroud P, Berland M, Dumas AM, Kennedy H (2002) Early specification of the hierarchical organization of visual cortical areas in the macaque monkey. Cerebral Cortex 12:453–465.

Bedny M (2017) Evidence from Blindness for a Cognitively Pluripotent Cortex. Trends in Cognitive Sciences 21:637–648.

Bedny M, Pascual-Leone A, Dodell-Feder D, Fedorenko E, Saxe R (2011) Language processing in the occipital cortex of congenitally blind adults. Proceedings of the National Academy of Sciences 108:4429–4434.

Bengtsson SL, Nagy Z, Skare S, Forsman L, Forssberg H, Ullén F (2005) Extensive piano practicing has regionally specific effects on white matter development. Nature Neuroscience 8:1148–1150.

Büchel C (2003) Cortical hierarchy turned on its head. Nature Neuroscience 6:657–658.

Burton H (2003) Visual cortex activity in early and late blind people. Journal of Neuroscience 23:4005–4011.

Butt OH, Benson NC, Datta R, Aguirre GK (2013) The fine-scale functional correlation of striate cortex in sighted and blind people. Journal of Neuroscience 33:16209–16219.

Chen X, Lu B, Yan CG (2018) Reproducibility of R-fMRI metrics on the impact of different strategies for multiple comparison correction and sample sizes. Human Brain Mapping 39:300–318.

Citri A, Malenka RC (2008) Synaptic plasticity: Multiple forms, functions, and mechanisms. Neuropsychopharmacology 33:18–41.

Cohen LG, Celnik P, Pascual-Leone A, Corwell B, Faiz L, Dambrosia J, Honda M, Sadatok N, Gerloff C, Catala MD, Hallett M (1997) Functional relevance of cross-modal plasticity in blind humans. Nature 389:180–183.

Collignon O, Dormal G, Albouy G, Vandewalle G, Voss P, Phillips C, Lepore F (2013) Impact of blindness onset on the functional organization and the connectivity of the occipital cortex. Brain 136:2769–2783.

Coppins N, Barlow-Brown F (2006) Reading difficulties in blind, braille-reading children. British Journal of Visual Impairment 24:37–39.

Cox RW (1996) AFNI: Software for analysis and visualization of functional magnetic resonance neuroimages. Computers and Biomedical Research 29:162–173.

Destrieux C, Fischl B, Dale A, Halgren E (2010) Automatic parcellation of human cortical gyri and sulci using standard anatomical nomenclature. NeuroImage 53:1–15.

Forkel SJ, Thiebaut de Schotten M, Kawadler JM, Dell’Acqua F, Danek A, Catani M (2014) The anatomy of fronto-occipital connections from early blunt dissections to contemporary tractography. Cortex 56:73–84.

Friston K (2010) The free-energy principle: A unified brain theory? Nature Reviews Neuroscience 11:127–138.

Friston KJ, Harrison L, Penny W (2003) Dynamic causal modelling. NeuroImage 19:1273–1302.

Fujii T, Tanabe HC, Kochiyama T, Sadato N (2009) An investigation of cross-modal plasticity of effective connectivity in the blind by dynamic causal modeling of functional MRI data. Neuroscience Research 65:175–186.

Hebb DO (1950) The Organization of Behavior; A Neuropsychological Theory. The American Journal of Psychology 63:633.

Heimler B, Amedi A (2020) Are critical periods reversible in the adult brain? Insights on cortical specializations based on sensory deprivation studies. Neuroscience and Biobehavioral Reviews 116:494–507.

Jiao Y, Lu X, Xu Y (2018) A graphical tactile display for the visually impaired. [C] Proceedings of AsiaHaptics, Best Video Demo Award.

Jung WH, Lee S, Lerman C, Kable JW (2018) Amygdala Functional and Structural Connectivity Predicts Individual Risk Tolerance. Neuron 98:394–404.e4.

Katarzyna Rączy, Urbańczyk A, Korczyk M, Szewczyk JM, Sumera E, Szwed M (2019) Orthographic Priming in Braille Reading as Evidence for Task-specific Reorganization in the Ventral Visual Cortex of the Congenitally Blind. Journal of Cognitive Neuroscience 31:1065–1078.

Keuroghlian AS, Knudsen EI (2007) Adaptive auditory plasticity in developing and adult animals. Progress in Neurobiology 82:109–121.

Kim JS, Kanjlia S, Merabet LB, Bedny M (2017) Development of the Visual Word Form Area Requires Visual Experience: Evidence from Blind Braille Readers. The Journal of Neuroscience 37:11495–11504.

Liu Y, Yu C, Liang M, Li J, Tian L, Zhou Y, Qin W, Li K, Jiang T (2007) Whole brain functional connectivity in the early blind. Brain 130:2085–2096.

Marshall L, Moys J-L (2020) Readers’ experiences of Braille in an evolving technological world. Visible Language 54.

Mason LK (2012) Experimental investigation of hand and finger usage in braille reading. Dissertations 209.

Matuszewski J, Kossowski B, Bola Ł, Banaszkiewicz A, Paplińska M, Gyger L, Kherif F, Szwed M, Frackowiak RS, Jednoróg K, Draganski B, Marchewka A (2021) Brain plasticity dynamics during tactile Braille learning in sighted subjects: Multi-contrast MRI approach. NeuroImage 227:117613.

Mongillo G, Barak O, Tsodyks M (2008) SynaptiC Theory of Working Memory. Science 319:1543–1546.

Noppeney U (2003) Effects of visual deprivation on the organization of the semantic system. Brain 126:1620–1627.

Pan S, Mayoral SR, Choi HS, Chan JR, Kheirbek MA (2020) Preservation of a remote fear memory requires new myelin formation. Nature Neuroscience 23.

Pascual-Leone A, Amedi A, Fregni F, Merabet LB (2005) the Plastic Human Brain Cortex. Annual Review of Neuroscience 28:377–401.

Pavani F, Röder B (2012) Crossmodal plasticity as a consequence of sensory loss: Insights from blindness and deafness. In: The new handbook of multisensory processes. (Stein BE, ed), pp 737–760. Cambridge, MA: MIT Press.

Penny WD, Stephan KE, Daunizeau J, Rosa MJ, Friston KJ, Schofield TM, Leff AP (2010) Comparing families of dynamic causal models. PLoS Computational Biology 6.

Power JD, Barnes KA, Snyder AZ, Schlaggar BL, Petersen SE (2012) Spurious but systematic correlations in functional connectivity MRI networks arise from subject motion. NeuroImage 59:2142–2154.

Reich L, Szwed M, Cohen L, Amedi A (2011) A ventral visual stream reading center independent of visual experience. Current Biology 21:363–368.

Ren K, Liu S, Lin M, Wang Y, Zhang QM (2008) A compact electroactive polymer actuator suitable for refreshable Braille display. Sensors and Actuators, A: Physical 143:335–342.

Ren Y, Nguyen VT, Sonkusare S, Lv J, Pang T, Guo L, Eickhoff SB, Breakspear M, Guo CC (2018) Effective connectivity of the anterior hippocampus predicts recollection confidence during natural memory retrieval. Nature Communications 9.

Renier L, De Volder AG, Rauschecker JP (2014) Cortical plasticity and preserved function in early blindness. Neuroscience and Biobehavioral Reviews 41:53–63.

Röder B, Kekunnaya R, Guerreiro MJS (2021) Neural mechanisms of visual sensitive periods in humans. Neuroscience and Biobehavioral Reviews 120:86–99.

Röder B, Stock O, Bien S, Neville H, Rösler F (2002) Speech processing activates visual cortex in congenitally blind humans. European Journal of Neuroscience 16:930–936.

Rohlf S, Habets B, von Frieling M, Röder B (2017) Infants are superior in implicit crossmodal learning and use other learning mechanisms than adults. eLife 6:1–23.

Ryles R (1996) The impact of braille reading skills on employment, income, education, and reading habits. Journal of Visual Impairment and Blindness 90:219–226.

Ryles R (2000) Braille as a predictor of success. In: Braille into the next millennium (Judith Dixon, ed), pp 463–491. Washington, DC: National Library Service for the Blind and Print Disabled, and Friends of Libraries for Blind and Physically Handicapped Individuals in North America.

Sabbah N, Authié CN, Sanda N, Mohand-Saïd S, Sahel JA, Safran AB, Habas C, Amedi A (2016) Increased functional connectivity between language and visually deprived areas in late and partial blindness. NeuroImage 136:162–173.

Sadato N (2005) How the blind “see” Braille: Lessons from functional magnetic resonance imaging. Neuroscientist 11:577–582.

Sadato N, Pascual-Leone A, Grafman J, Deiber MP, Ibañez V, Hallett M (1998) Neural networks for Braille reading by the blind. Brain 121:1213–1229.

Schepers IM, Hipp JF, Schneider TR, Röder B, Engel AK (2012) Functionally specific oscillatory activity correlates between visual and auditory cortex in the blind. Brain 135:922–934.

Scholz J, Klein MC, Behrens TEJ, Johansen-Berg H (2009) Training induces changes in white-matter architecture. Nature Neuroscience 12:1370–1371.

Schroeder FK (1996) Perceptions of Braille Usage by Legally Blind Adults. Journal of Visual Impairment & Blindness 90:210–218.

Seghier ML, Josse G, Leff AP, Price CJ (2011) Lateralization is predicted by reduced coupling from the left to right prefrontal cortex during semantic decisions on written words. Cerebral Cortex 21:1519–1531.

Striem-Amit E, Ovadia-Caro S, Caramazza A, Margulies DS, Villringer A, Amedi A (2015) Functional connectivity of visual cortex in the blind follows retinotopic organization principles. Brain 138:1679–1695.

Tingley D, Yamamoto T, Hirose K, Keele L, Imai K (2014) Mediation: R package for causal mediation analysis. Journal of Statistical Software 59:1–38.

Trent SD, Truan MB (1997) Speed, accuracy, and comprehension of adolescent braille readers in a specialized school. Journal of Visual Impairment and Blindness 91:494–500.

Völkel T, Weber G, Baumann U (2008) Tactile graphics revised: The novel BrailleDis 9000 pin-matrix device with multitouch input. Lecture Notes in Computer Science (including subseries Lecture Notes in Artificial Intelligence and Lecture Notes in Bioinformatics) 5105 LNCS:835–842.

Wang X, Caramazza A, Peelen M V., Han Z, Bi Y (2015) Reading without speech sounds: VWFA and its connectivity in the congenitally deaf. Cerebral Cortex.

Watkins KE, Cowey A, Alexander I, Filippini N, Kennedy JM, Smith SM, Ragge N, Bridge H (2012) Language networks in anophthalmia: Maintained hierarchy of processing in “visual” cortex. Brain 135:1566–1577.

Winkler AM, Ridgway GR, Douaud G, Nichols TE, Smith SM (2016) Faster permutation inference in brain imaging. NeuroImage 141:502–516 Available at: http://dx.doi.org/10.1016/j.neuroimage.2016.05.068.

Wittenberg GF, Werhahn KJ, Wassermann EM, Herscovitch P, Cohen LG (2004) Functional connectivity between somatosensory and visual cortex in early blind humans. European Journal of Neuroscience 20:1923–1927.

Yan CG, Cheung B, Kelly C, Colcombe S, Craddock RC, Di Martino A, Li Q, Zuo XN, Castellanos FX, Milham MP (2013) A comprehensive assessment of regional variation in the impact of head micromovements on functional connectomics. NeuroImage 76:183–201.

Yan CG, Wang X Di, Zuo XN, Zang YF (2016) DPABI: Data Processing & Analysis for (Resting-State) Brain Imaging. Neuroinformatics 14:339–351.

Zeidman P, Jafarian A, Corbin N, Seghier ML, Razi A, Price CJ, Friston KJ (2019) A guide to group effective connectivity analysis, part 1: First level analysis with DCM for fMRI. NeuroImage 200:174–190.

