## Supplementary Information for "Two tales of one neural link predict blind individual’s Braille reading proficiency"

**Supplementary Figure S1. Related to Figure 2.**

Functional connectivity patterns for the right primary 'visual' cortex (V1) and the right lateral occipital cortex (LOC).

**Supplementary Figure S2. Related to Figure 2.**

Enhanced functional connectivity between the left lateral occipital cortex (LOC) and the left intraparietal sulcus (IPS) in the late blind individuals compared to sighted controls.

**Supplementary Figure S3. Related to Figure 3.**

Correlations between the 'visual'-frontal functional connectivity and the Braille reading practice time in childhood and adulthood of late blind individuals respectively.

**Supplementary Figure S4. Related to Figure 4.**

Experimental design and brain activations across blind subjects during the natural Braille reading task.

**Supplementary Figure S5. Related to Figure 5.**

Relationships between Braille reading task modulation of the LOC-IFC effective connectivity and Braille reading proficiency of late blind individuals.

**Supplementary Table S1. Related to Figure 4.**

Endogenous connectivity parameters (in Hz) from the BMA analysis of the late blind group.

**Supplementary Table S2. Related to Figure 4.**

Modulatory effect parameters (in Hz) from the BMA analysis of the late blind group.

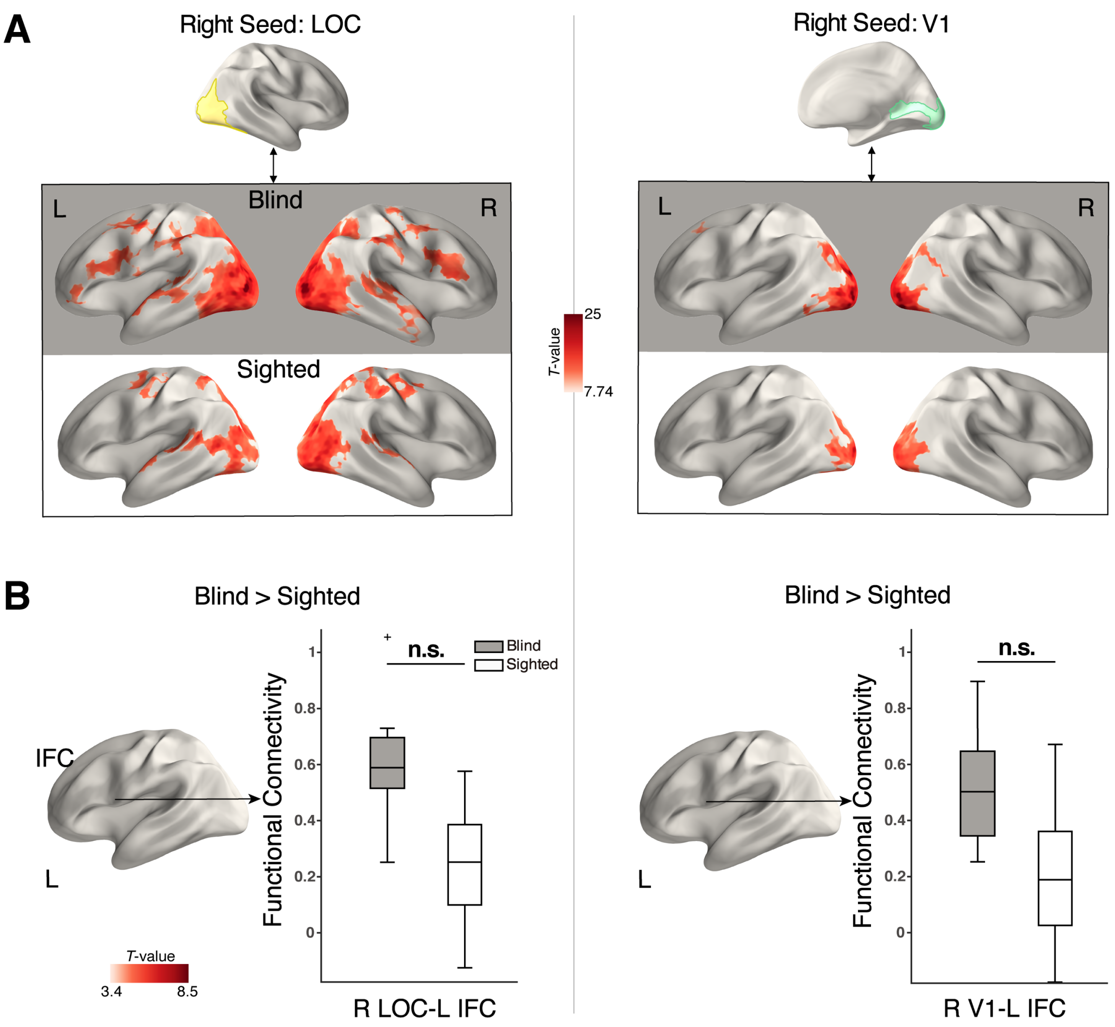


**Supplementary Figure S1. Related to Figure 2.** Functional connectivity patterns for the right primary 'visual' cortex (V1) and the right lateral occipital cortex (LOC). (A) Maps show vertices with significantly positive connectivity with the right LOC seed (vertexwise $p < 10^{-6}$, uncorrected, cluster size > 100 mm) in the late blind group and sighted group (left panel). Maps show vertices with significant positive connectivity with the right V1 seed (vertexwise $p < 10^{-6}$, uncorrected, cluster size > 100 mm) in the late blind group and sighted group (right panel). (B) No significant vertices were found in functional connectivity patterns for right LOC (left panel) and V1 (right panel) of the late blind group versus the sighted group, respectively ($p < 0.01$, corrected for both hemispheres, cluster size > 30 mm). Box plot of connectivity strengths (Fisher r-to-z-transformed)

between the right seed and the left IFC cluster (located from the left panel in Figure 2B) for the two groups. The horizontal line denotes the median, and the asterisk indicates significance. V1 primary visual cortex (light green), LOC lateral occipital cortex (light yellow), IFC inferior frontal cortex. L left hemisphere, R right hemisphere. Blind, dark color; Sighted, light color. ** $p < 0.01$, corrected for both hemispheres. n.s. non-significant.

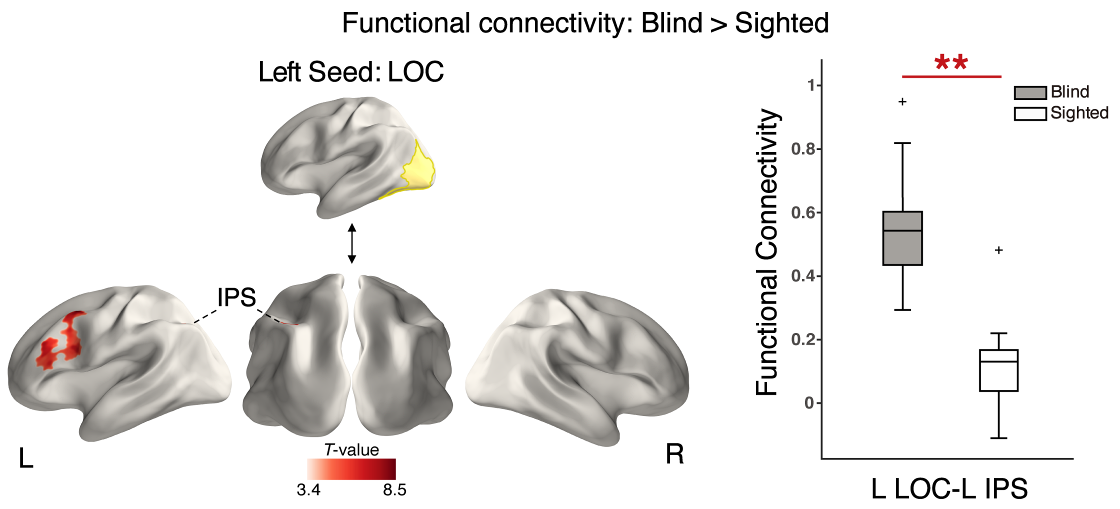


**Supplementary Figure S2. Related to Figure 2.** Enhanced functional connectivity between the left lateral occipital cortex (LOC) and the left intraparietal sulcus (IPS) in late blind individuals compared to sighted controls. Significant vertices were corrected for multiple comparisons using permutation testing with threshold-free cluster enhancement (two-sample *T*-test, $p < 0.01$, corrected for both hemispheres, cluster size > 30 mm). Box plot of connectivity strengths (Fisher r-to-z-transformed) between the left LOC and the left IPS cluster for the two groups. The horizontal line denotes the median, and the asterisk indicates significance. LOC lateral occipital cortex (light yellow), IPS intraparietal sulcus. L left hemisphere, R right hemisphere. Blind, dark color; Sighted, light color.

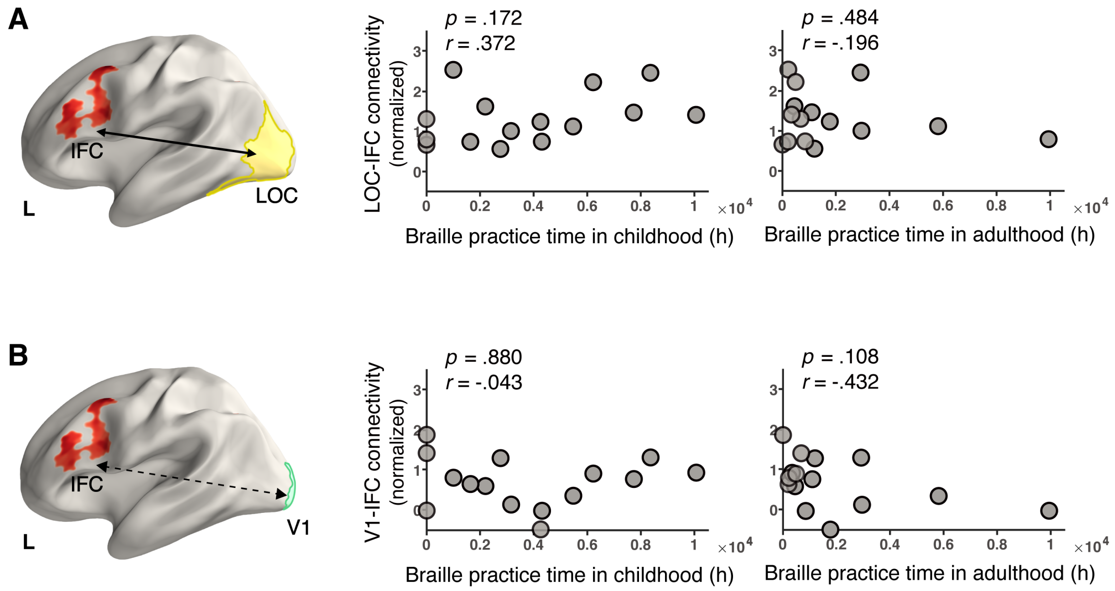


**Supplementary Figure S3. Related to Figure 3.** Correlations between the ‘visual’-frontal functional connectivity and the Braille reading practice time in childhood and adulthood of late blind individuals, respectively. L left hemisphere. All *r* and *p*-value pairs reflect Pearson correlation, consistent with the Spearman correlation results.

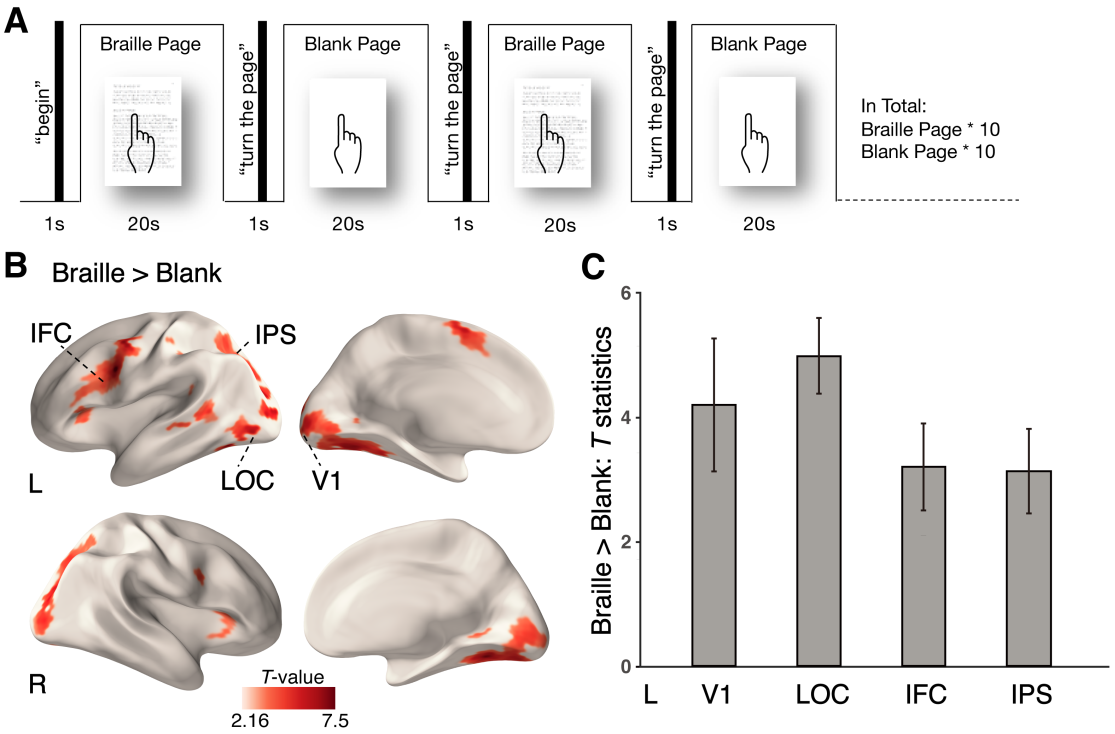


**Supplementary Figure S4. Related to Figure 4.** Experimental design and brain activations across blind subjects during the natural Braille reading task. (A) Experiment paradigm including natural Braille reading pages and blank counterpart. And examples of natural Braille articles. (B) Activated brain regions revealed by the Braille > blank contrast, mainly including bilateral V1, LOC, IPS, and left-lateralized IFC. Significant activations were identified using a vertexwise threshold ($p < 0.025$ for both hemispheres, uncorrected, cluster size > 2 vertices). (C) Average *T* statistics in left V1, LOC, IFC, and IPS in the late blind group. Error bars signify SEM. L left hemisphere, R right hemisphere.

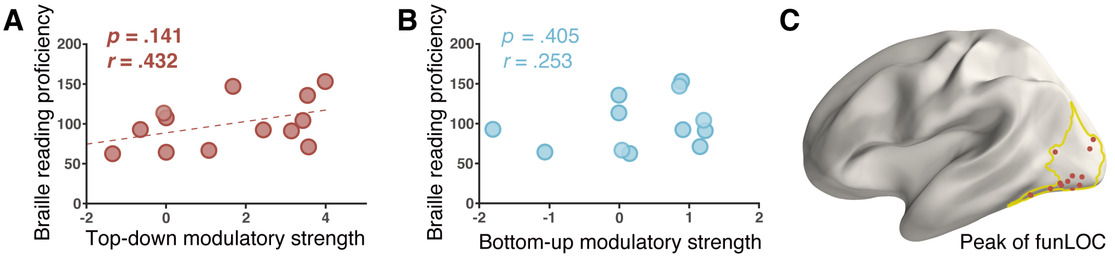


**Supplementary Figure S5. Related to Figure 5.** Relationships between Braille reading task modulation of the LOC-IFC effective connectivity and Braille reading proficiency of late blind individuals. (A) The positive relationship between the top-down modulation of the left IFC to LOC connectivity and Braille reading proficiency tended to approach significance. (B) The bottom-up modulation of the left LOC to IFC connectivity was not significantly correlated with Braille reading proficiency. (C) Red dots indicated individual peak vertices of functional LOC during the natural Braille reading task, which acted as the neural mediator of the top-down modulation in support of Braille reading proficiency. Anatomical LOC boundary (light yellow). All *r* and *p*-value pairs reflect Pearson correlation.

**Supplementary Table S1. Related to Figure 4.** Endogenous connectivity parameters (in Hz) from the BMA analysis of the late blind group.

| Effective connectivity | Mean | SD |
|---|---|---|
| IPS → LOC | 0.4069 | 0.0241 |
| LOC → IPS | -0.4068 | 0.0255 |
| LOC → IFC | 0.1342 | 0.0144 |
| IFC → LOC | -0.3917 | 0.0265 |
| LOC → V1 | 0.5759 | 0.0167 |
| V1 → LOC | -0.1259 | 0.0247 |

**Supplementary Table S2. Related to Figure 4.** Modulatory effect parameters (in Hz) from the BMA analysis of the late blind group.

| Modulatory effect | Mean | SD |
|---|---|---|
| IPS → LOC | 1.0618 | 0.0949 |
| LOC → IPS | -1.5590 | 0.0938 |
| LOC → IFC | 0.5474 | 0.0481 |
| IFC → LOC | 1.4958 | 0.1268 |
